# High-quality genome assemblies uncover caste-specific long non-coding RNAs in ants

**DOI:** 10.1101/155119

**Authors:** Emily J. Shields, Roberto Bonasio

## Abstract

Ants are an emerging model system for neuroepigenetics, as embryos with virtually identical genomes develop into different adult castes that display strikingly different physiology, morphology, and behavior. Although a number of ant genomes have been sequenced to date, their draft quality is an obstacle to sophisticated analyses of epigenetic gene regulation. Using long reads generated with Pacific Biosystem single molecule real time sequencing, we have reassembled *de novo* high-quality genomes for two ant species: *Camponotus floridanus* and *Harpegnathos saltator*. The long reads allowed us to span large repetitive regions and join sequences previously found in separate scaffolds, leading to comprehensive and accurate protein-coding annotations that facilitated the identification of a *Gp-9-like* gene as differentially expressed in *Harpegnathos* castes. The new assemblies also enabled us to annotate long non-coding RNAs for the first time in ants, revealing several that were specifically expressed during *Harpegnathos* development and in the brains of different castes. These upgraded genomes, along with the new coding and non-coding annotations, will aid future efforts to identify epigenetic mechanisms of phenotypic and behavioral plasticity in ants.

## INTRODUCTION

Social insects are of great interest to epigenetics because they display remarkable phenotypic plasticity within the boundaries of a single genome^1,2^. Among social insects, the ponerine ant *Harpegnathos saltator* is emerging as a model system to study the epigenetic regulation of brain function and behavior. *Harpegnathos* workers have the ability to convert to acting queens, called gamergates, that are allowed to mate and lay fertilized eggs. This transition allows genetic access to the germline, similar to non-social model organisms^3^. We have previously shown that the conversion of workers to gamergates is accompanied by major changes in gene expression in their brains^4^, but the epigenetic mechanisms responsible for these changes remain largely unknown.

Previous work in *Harpegnathos* and a more conventional ant species, the Florida carpenter ant *Camponotus floridanus*, has suggested that epigenetic pathways, including those that control histone modifications and DNA methylation, might be responsible for differential deployment of caste-specific traits^5–7^. In fact, pharmacological and molecular manipulation of histone acetylation affects caste-specific behavior in *Camponotus*^8^, suggesting a direct role for epigenetics in the social behavior of these ants.

Although the molecular mechanisms by which environmental and developmental cues are converted into epigenetic information on chromatin remain subject of intense investigation^9^, it has become increasingly clear that non-coding RNAs play an important role in mediating this flow of information^10^. In particular, long non-coding RNAs (lncRNAs)—transcripts longer than 200 bp that are not translated into proteins—have been proposed to function as a communication conduit between the genome and chromatin-associated complexes^11^, thus participating in the epigenetic regulation of gene expression^12,13^. Some envision a model in which lncRNAs recruit machinery that mediates epigenetic silencing, such as CoREST^14^, Polycomb group proteins^15–19^ and DNA methyltransferases^20^, or epigenetic activation, such as the MLL complex^21^. LncRNAs have also been implicated in maintaining looping interactions from promoters to enhancers^22^, and as organizers of 3D genome architecture^23^.

LncRNAs have been annotated extensively in human^24,25^, mouse^26,27^, other model organisms including zebrafish, *Drosophila melanogaster* and *Caenorhabditis elegans*^28–32^, and the honeybees *Apis mellifera* and *Apis cerana*^33^, but to our knowledge no comprehensive annotation of lncRNAs in an ant species has been reported. This is in part because ant genomes, including those of *Camponotus* and *Harpegnathos*^5^, are still in low-quality draft form due to the prevalent use of whole genome shotgun sequencing to assemble them. In addition to making lncRNA annotation practically impossible, these low-quality genomes also hamper the sophisticated genome-wide analyses required for epigenetic research, thus limiting the reach of these species as model organisms.

We upgraded the genomes of *Harpegnathos* and *Camponotus* to megabase-level with a combination of *de novo* assembly of Pacific Biosystems (PacBio) long reads, scaffolding with mate pairs and long reads, and polishing with short reads. The contiguity of both genomes (measured by the number of contigs) improved by at least 24-fold while maintaining the high accuracy of the short-read only assemblies. The new assemblies were used to annotate protein-coding genes and lncRNAs, leading to the discovery of lncRNAs differentially expressed between *Harpegnathos* castes and developmental stages. We anticipate that these improvements to the *Harpegnathos* and *Camponotus* genomes will lead to greater understanding of the genetic and epigenetic factors that underlie the behavior of these social insects.

## RESULTS

### Long-read sequencing yields highly contiguous assemblies

We sequenced genomic DNA isolated from *Harpegnathos* and *Camponotus* workers using PacBio single molecule real time (SMRT) technology, which produced reads of insert (ROI) with median sizes of 7.5 kb and 10 kb for *Harpegnathos* and *Camponotus*, respectively (**Fig. S1**). These reads are much longer than those used for whole-genome shotgun draft assemblies, including the previously reported assemblies for these two ant species^5^, and are thus expected to yield longer contigs and scaffolds with fewer gaps (scheme, **Fig. 1A**). After extracting ROI from the raw PacBio reads we obtained a total sequence coverage of 70x for *Harpegnathos* and 53x for *Camponotus*, compatible with PacBio-only genome assembly^34^.

**Figure 1.**
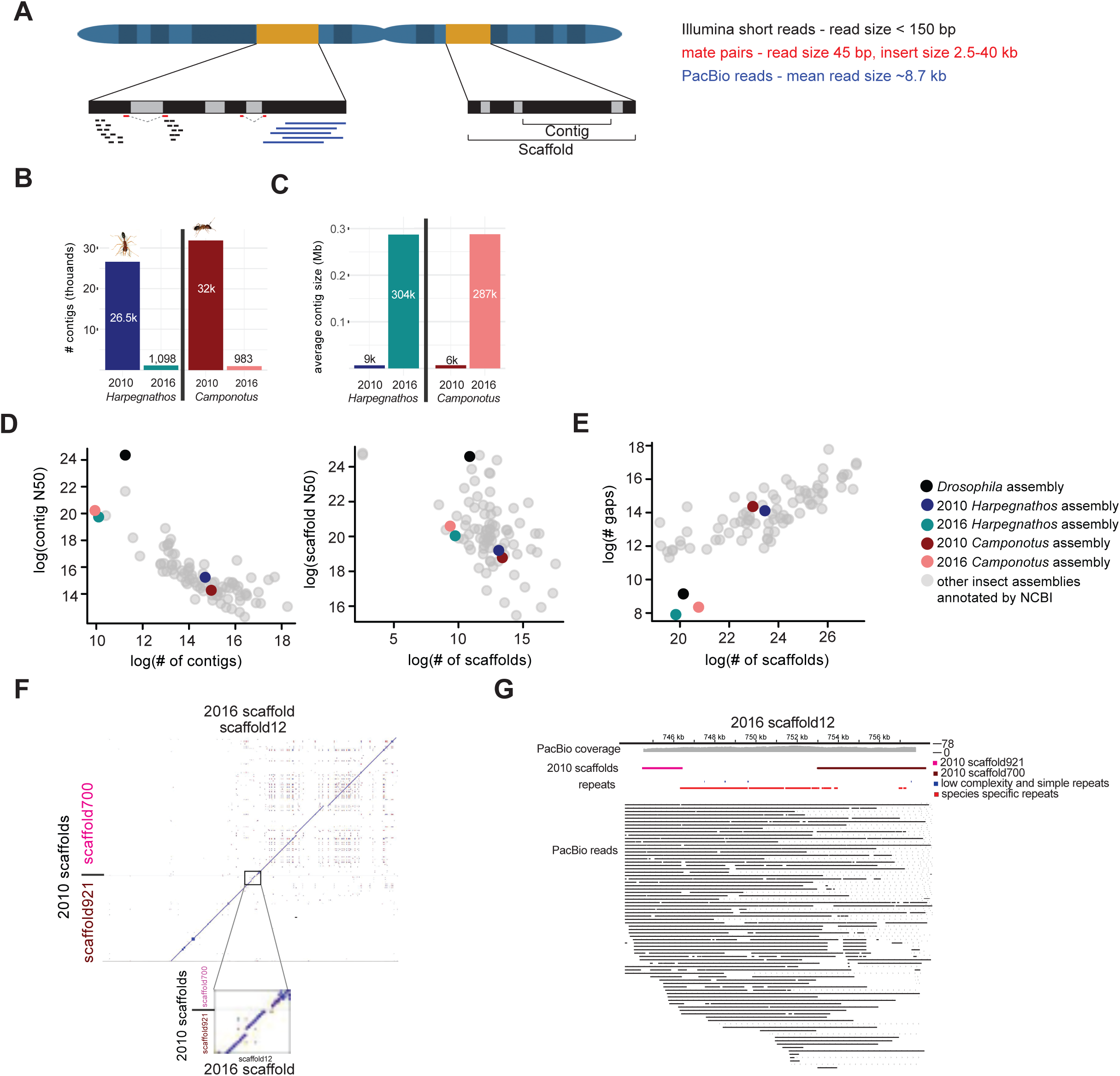
PacBio sequencing improves assemblies for two ant genomes. **(A)** Scheme showing types of reads used in assembly. PacBio SMRT sequencing (“PacBio reads,” blue) is a third-generation sequencing technology that produces reads as long as 70 kb, with a mean size of ∼8.7 kb. These reads were used in the initial *de novo* assembly. Mate pairs (red) are generated from sequencing the circularized and ligated ends of long insert libraries yielding short reads from two regions of the genome separated by the insert size (ranging from 2.5 kb to 40 kb), and aided in scaffolding the assemblies. Illumina short reads (black) are < 150 bp but have a lower sequencing error rate than PacBio sequenced reads, and were used to correct sequence errors in the assembly. **(B)** 2016 *Harpegnathos* and *Camponotus* genome assemblies have a greatly reduced number of contigs compared to the 2010 assemblies. **(C)** The average contig size is higher in the 2016 assemblies compared to the 2010 assemblies. **(D)** Comparison of *Harpegnathos* and *Camponotus* genome assemblies to other insect genomes using contig number and N50 (left) and scaffold number and N50 (right). The N50 is a measure of genome quality, and is defined as the length of the contig or scaffold for which the summed lengths of all contigs/scaffolds of the size or larger equal at least half the genome size. All scaffold-level insect assemblies annotated by NCBI as of 5/8/17 are included in the comparison, with the *Drosophila* assembly (black), 2010 *Harpegnathos* and *Camponotus* assemblies (2010 *Harpegnathos*, maroon; 2010 *Camponotus*, dark blue), and 2016 *Harpegnathos* and *Camponotus* assemblies (2016 *Harpegnathos*, teal; 2016 *Camponotus*, coral) highlighted. **(E)** Number of gaps and gapped bases in insect assemblies, with the same insects included as in **(D)**. **(F)** Two separate scaffolds from the 2010 *Harpegnathos* assembly map to the same 2016 scaffold. The 2010 scaffolds, scaffold921 and scaffold700, are depicted along the y-axis, with the 2016 scaffold, scaffold12, along the x-axis. Dots indicate regions where there is significant sequence similarity. The boundary region between the 2010 scaffolds is shown in the inset. **(G)** A genome browser view of region from **(F)** shows coverage by several PacBio reads that span the stretch of repetitive sequence across the gap between the two 2010 scaffolds.

We used these long reads to assemble the two genomes *de novo* using a multi-step process (**Fig. S2A**). The initial *de novo* assembly was performed with the dedicated long read assembler Canu^34^, followed by polishing with Quiver^35^. Although these initial steps produced assemblies that surpassed the contiguity of the current draft genomes (**Fig. S2A**, **Table 1**), we leveraged previously generated sequencing data to maximize the quality of the newly assembled genomes. Scaffolding using PBJelly^35^ and SSPACE-standard^36^ was followed by error correction using short reads with Pilon^37^, which takes advantage of the higher accuracy of the Illumina platform.

**Table 1.** Genome quality metrics for old and new assemblies

|  | H. sal |  | C. flo |  |
| --- | --- | --- | --- | --- |
|  | 2010 assembly | 2016 assembly | 2010 assembly | 2016 assembly |
| number of contigs | 26,592 | 1,098 | 31,883 | 983 |
| contig N50 | 39,378 | 875,847 | 18,762 | 1,225,609 |
| number of scaffolds | 8,893 | 858 | 10,791 | 657 |
| scaffold N50 (bp) | 601,965 | 1,078,644 | 451,320 | 1,585,631 |
| longest scaffold (bp) | 2,276,656 | 3,353,128 | 2,671,896 | 10,163,455 |
| number of gaps | 17,699 | 240 | 21,092 | 326 |
| number of Ns | 11,466,753 | 933,241 | 8,173,001 | 1,771,909 |
| total size (bp) | 294,465,601 | 335,266,283 | 232,685,334 | 284,009,204 |

The new PacBio sequencing-derived assemblies (“2016 assemblies”) compared favorably to the short-read assemblies currently available for both ant species (“2010 assemblies”). Despite capturing a larger amount of genomic sequence (**Table 1**), the number of contigs was dramatically decreased in the 2016 assemblies (**Fig. 1B**) and, consequently, their average size was more than 30-fold larger than in the 2010 assemblies (**Fig. 1C**), reflecting greatly increased assembly contiguity. Scaffolding was also improved in the 2016 assemblies, which consisted of fewer scaffolds that were larger (**Fig. S2B**) and contained fewer gaps than the 2010 assemblies (**Table 1**). Improvements were also evident in the conventional metrics of assembly quality such as contig and scaffold N50s (**Table 1**). Overall the contig N50 size grew 22-fold and 65-fold larger for *Harpegnathos* and *Camponotus* respectively and in both assemblies the scaffold N50 size surpassed 1 Mb (**Table 1**).

The N50 contig size of our improved *Harpegnathos* and *Camponotus* assemblies top almost all other insect genomes available in the NCBI database, with the exception of two genomes also assembled using PacBio long read sequencing (*Drosophila serrata*^38^ and *Aedes albopictus*) and the classic model organism *Drosophila melanogaster* (**Fig. 1D**, left). The number and size of scaffolds also compared favorably with other available genomes (**Fig. 1D,** right), and, most notably, the number of gaps in our new assemblies was lower than for any other insect genome in this set, including *Drosophila melanogaster* (**Fig. 1E**).

PacBio reads can span long repetitive sequence that cannot be assembled properly by whole-genome shotgun sequencing using short reads^39^. We found several cases where distinct scaffolds from the 2010 assemblies mapped to a single new scaffold (or contig) in the 2016 assemblies, separated by repetitive sequences. For example, scaffolds 921 and 700 from 2010 were joined as contiguous parts of a larger scaffold in the improved 2016 assemblies (**Fig. 1F**), separated by ∼6.5 kb of repeats that were spanned by multiple PacBio reads (**Fig. 1G**). Indeed, much of the new assembled DNA sequence that was missing from the 2010 assemblies consisted of repeats (**Fig. S3A**), largely species-specific repeats, with some contribution from retroelements and DNA transposons (**Fig. S3B**).

Thus, the much longer reads obtained with PacBio sequencing allowed us to assemble across longer repeats than previously possible, resulting in updated *Harpegnathos* and *Camponotus* genomes with better contiguity than most insect genomes available at the time of writing.

### Long-read assemblies are highly accurate

The major drawback of PacBio sequencing is its high error rate, estimated to be 10-15%, compared to the Illumina short-read sequencing error rate of ∼0.2-0.8%^40^. We countered this limitation with deep sequence coverage (70x and 53x, see above) and by polishing our assemblies with the large amount of Illumina short-read sequences generated for the original draft genomes^5^. Nonetheless, we wished to determine that the improved assembly contiguity did not come at the expense of sequence accuracy. One relevant metric for genome quality with practical consequences for gene expression measurements is the rate at which RNA-seq reads map to the assembly, with the caveat that this reports only on sequence accuracy at transcribed regions. We sequenced RNA from different developmental stages in both species and found that in all cases a significantly higher percentage of the reads mapped to the 2016 *Harpegnathos* and *Camponotus* assemblies compared to the 2010 draft versions (**Fig. 2A**). As these reads were not used to produce the assemblies, they provided an orthogonal method of evaluating genome completeness and accuracy. The improved mapping rate suggests that the new assemblies capture transcribed, previously unassembled sequence. Moreover, the mismatch rate per base was decreased (**Fig. 2B**), which demonstrates that our strategy to correct PacBio sequencing errors successfully generated highly accurate genome sequences.

**Figure 2.**
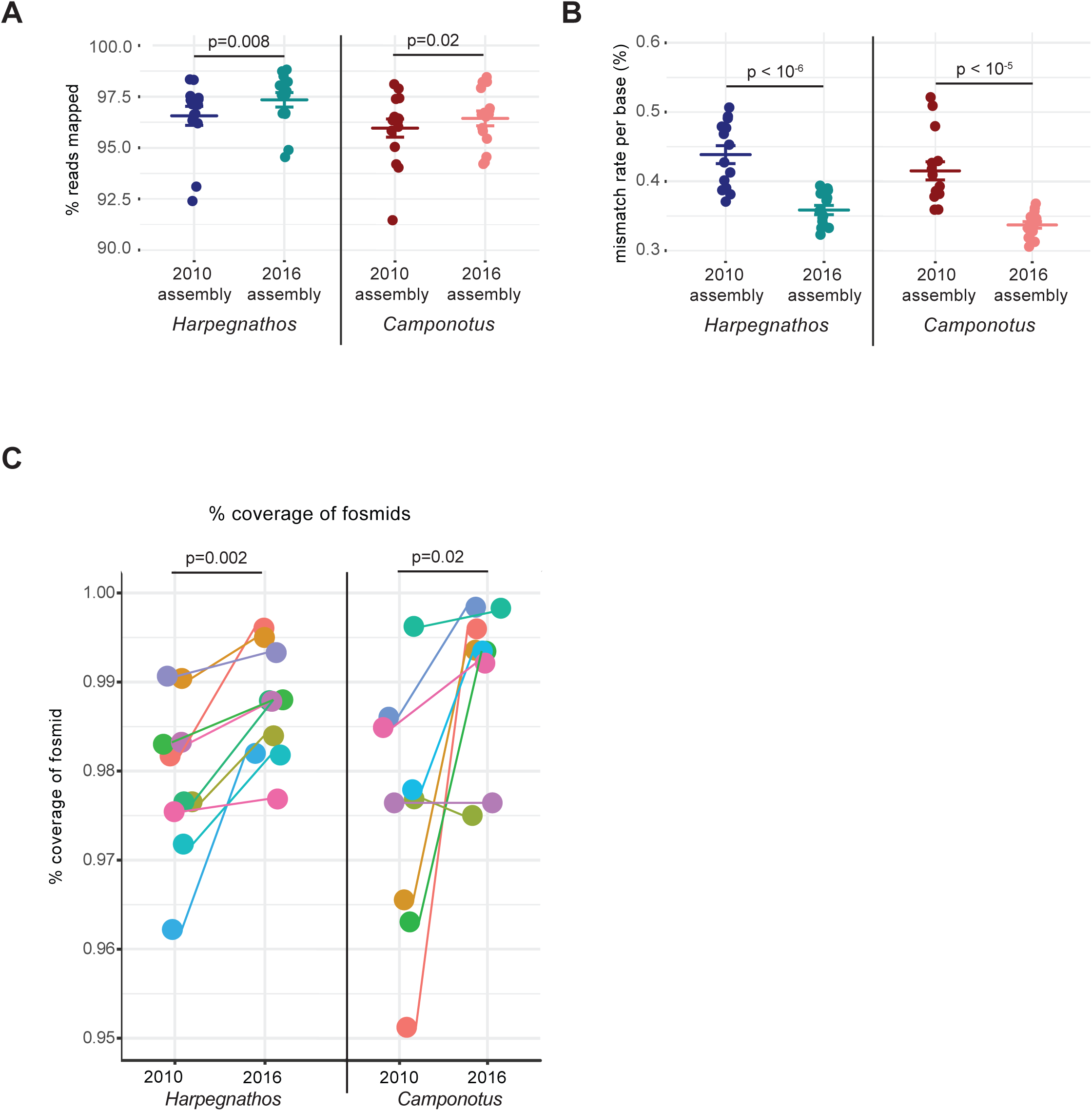
Improved accuracy of new assemblies. **(A–B)** Mapping (A) and sequence mismatch (B) rates for RNA-seq reads from various developmental stages of *Harpegnathos* (n=14) and *Camponotus* (n=15) to old and new assemblies. Horizontal bars indicate the means. P-values are from two-sided, paired Student’s t-test. **(C)** 2010 and 2016 assembly accuracy measured by % of fosmid Sanger sequence covered on a single scaffold. The scaffold with the highest similarity to the fosmid was found using BLAST, then a global alignment of the fosmid with that scaffold was performed. The % covered is calculated from the number of bases on the fosmid matching the scaffold. Each dot represents a fosmid. P-value is from a two-sided Student’s t-test.

To obtain an independent assessment of sequence and assembly accuracy that was not skewed toward transcribed regions, we analyzed the Sanger sequences of 10 (*Harpegnathos*) or 9 (*Camponotus*) fosmid clones of ∼40 kb that were previously generated to validate the short-read assemblies^5^, but were not used in the construction of either the 2010 or 2016 genomes. Alignment of these highly confident long sequences to the genomes showed similar or higher coverage in the new assemblies compared to the draft 2010 versions (**Fig. 2C**, **Table S1**).

### Protein-coding annotation captures new gene models

With improved genomes in hand, we sought to annotate protein-coding genes using a combination of *ab initio* transcriptome reconstruction of RNA-seq reads, homology-based searches with sequences from related organisms, and *de novo* identification of gene structures based on sequence features (**Fig. S4A**). We used the MAKER2^41^ pipeline to combine these sources of evidence, and retained gene models using both the annotation edit distance (**Fig. S4B**; AED^42^, a metric of agreement between evidence types) and the presence of proteins domains, measured by querying the protein families domains database (PFAM) maintained by EMBL^43^. Specifically, we removed from the annotation gene models that were only supported by one type of evidence (i.e. AED=1) and did not contain any discernible protein domains. We obtained final sets of 20,659 and 18,620 protein-coding genes for *Harpegnathos* and *Camponotus* respectively (**Fig. S4C**). Most of the gene models removed by this filtering step did not have any homology to other organisms, in addition to their lack of a PFAM domain, suggesting that they were spurious annotation products and did not correspond to real protein-coding genes (**Fig. S5**).

The filtered protein-coding annotations for *Harpegnathos* and *Camponotus* were evaluated for completeness against a core set of 1,066 evolutionarily conserved arthropod genes^44^. The new 2016 annotations recovered a slightly higher percentage of these core conserved genes compared to the 2010 annotations (**Table 2**).

**Table 2.** Quality metrics for protein-coding annotation

|  | H. sal |  | C. flo |  |
| --- | --- | --- | --- | --- |
|  | 2010 assembly | 2016 assembly | 2010 assembly | 2016 assembly |
| # genes in annotation | 18,564 | 20,659 | 17,064 | 18,620 |
| BUSCO results |  |  |  |  |
| complete | 98.4% | 98.6% | 97.2% | 98.1% |
| incomplete or missing | 1.6% | 1.4% | 2.8% | 1.9% |

Looking at a more comprehensive set of other genomes we found that the number of gene models encoding proteins conserved throughout evolution was more or less unchanged after the genome update (**Fig. 3A**). Interestingly, a higher percentage of genes in the 2016 assemblies had no homology to known protein-coding genes in human, mouse, and a panel of insects, including several Hymenoptera, all from annotations curated by NCBI (**Fig. 3B**, red boxes). However, a majority of these gene models without homology to known proteins contained at least one recognizable PFAM domain (**Fig. 3B**), suggesting that they might encode true protein-coding genes that might have been missed from previous annotation efforts in other organisms.

**Figure 3.**
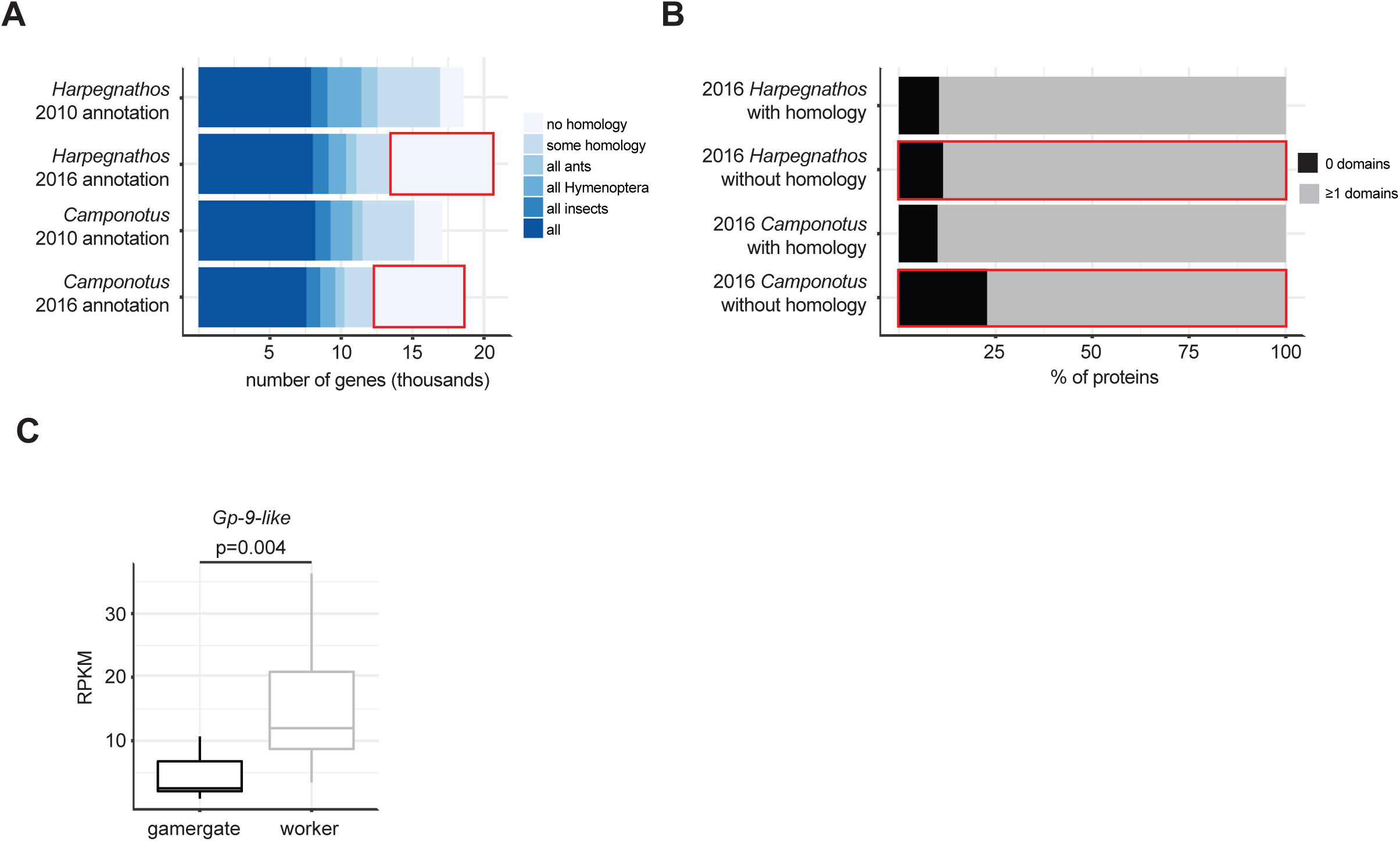
Annotation of protein-coding genes. **(A)** Number of genes in 2010 and 2016 *Harpegnathos* and *Camponotus* annotations with a homolog in a panel of other ants, Hymenoptera, and animals. A gene was considered ant-specific if it had a homolog in >90% of ant genomes available on NCBI and Hymenoptera-specific if it had a homolog in >90% of Hymenoptera genomes. The “all insects” category indicates genes with a homolog also in *Drosophila*, and the “all” category contains genes with homology to all insects and also a mammal (*Homo sapiens* or *Mus musculus*). **(B)** Fraction of genes with no detectable homology (outlined in red in **(A)**) that contains no (black) or more than 1 (gray) PFAM domains. **(C)** Expression of the newly annotated *Gp-9-like* gene in *Harpegnathos* gamergates (n=12) and workers (n=11). P-value is from a two-sided student’s t-test.

We reasoned that the improved assemblies and protein-coding annotations might uncover biologically relevant genes missing in the older versions. *Harpegnathos* workers are characterized by their unique reproductive and brain plasticity that, in absence of a queen, allows some of them to transition to a queen-like status accompanied by dramatic changes in physiology and behavior^5,6^. The converted workers are referred to as “gamergates”. We previously showed that this behavioral transition is accompanied by major changes in brain gene expression^4^. Reanalyzing this data set, we found that a *Gp-9-like* gene missing in the old annotations had significantly higher expression in worker brains compared to gamergates (**Fig. 3C**). This gene encodes one of several proteins with strong homology to a pheromone-binding protein well studied in the fire ant *Solenopsis invicta* because it marks a genomic element that governs colony structure^45^. A polymorphism in *S. invicta Gp-9* segregates with the ability of the colony to accept one fertile queen or several. Other ant species also including *Vollenhovia emeryi* and *Dinoponera quadriceps* have several genes encoding *Gp-9-like* homologs, some of which display caste-specific expression patterns (**Fig. S6**). Notably, the *Gp-9-like* homolog upregulated in *Dinoponera* worker brains is orthologous to the differentially expressed *Gp-9-like* gene in *Harpegnathos*, suggesting that the role of this pheromone-binding protein in social organization is more conserved than previously appreciated.

One specific locus where contiguity increases and improvements are made to the protein-coding annotation is the *Hox* cluster, a group of developmental genes with orthologs in many organisms^46^. As originally noted by Simola *et al*.^47^, homologs for two *Drosophila Hox* cluster genes, *lab* and *abd-A*, were surprisingly missing from the 2010 *Harpegnathos* annotation. However, both genes were correctly annotated in the new *Harpegnathos* genome and were properly positioned in the *Hox* cluster, in the same order as the corresponding *Drosophila* homologs (**Fig. 4A**). At a closer look, the 2010 annotation contained gene models overlapping the loci for *Iab* and *abd-A* but they were truncated, covering only 33% and 55% of the 2016 models (**Fig. 4B**, **C**, and data not shown), which had previously prevented their detection by homology searches. Importantly, the contiguity of the *Hox* cluster is critical to its function, as genes in the cluster are expressed in a collinear fashion during development and maintain the identity of different body segments^48^. *Drosophila* and the silkworm *Bombyx mori* have split *Hox* clusters^49,50^, but many other insects have an intact one^51–54^, including the fellow Hymenopteran *Apis mellifera*^55^. In our previous assemblies, the *Harpegnathos Hox* cluster was entirely contained in a single scaffold but the *Camponotus* cluster was split among three different clusters, begging the question of whether this separation was due to the actual relocalization of genes during evolution or simply discontinuous assembly. The improved 2016 assemblies answered this question by showing that the entire *Hox* clusters could be assembled into a single scaffold also in *Camponotus* (**Fig. 4A**).

**Figure 4.**
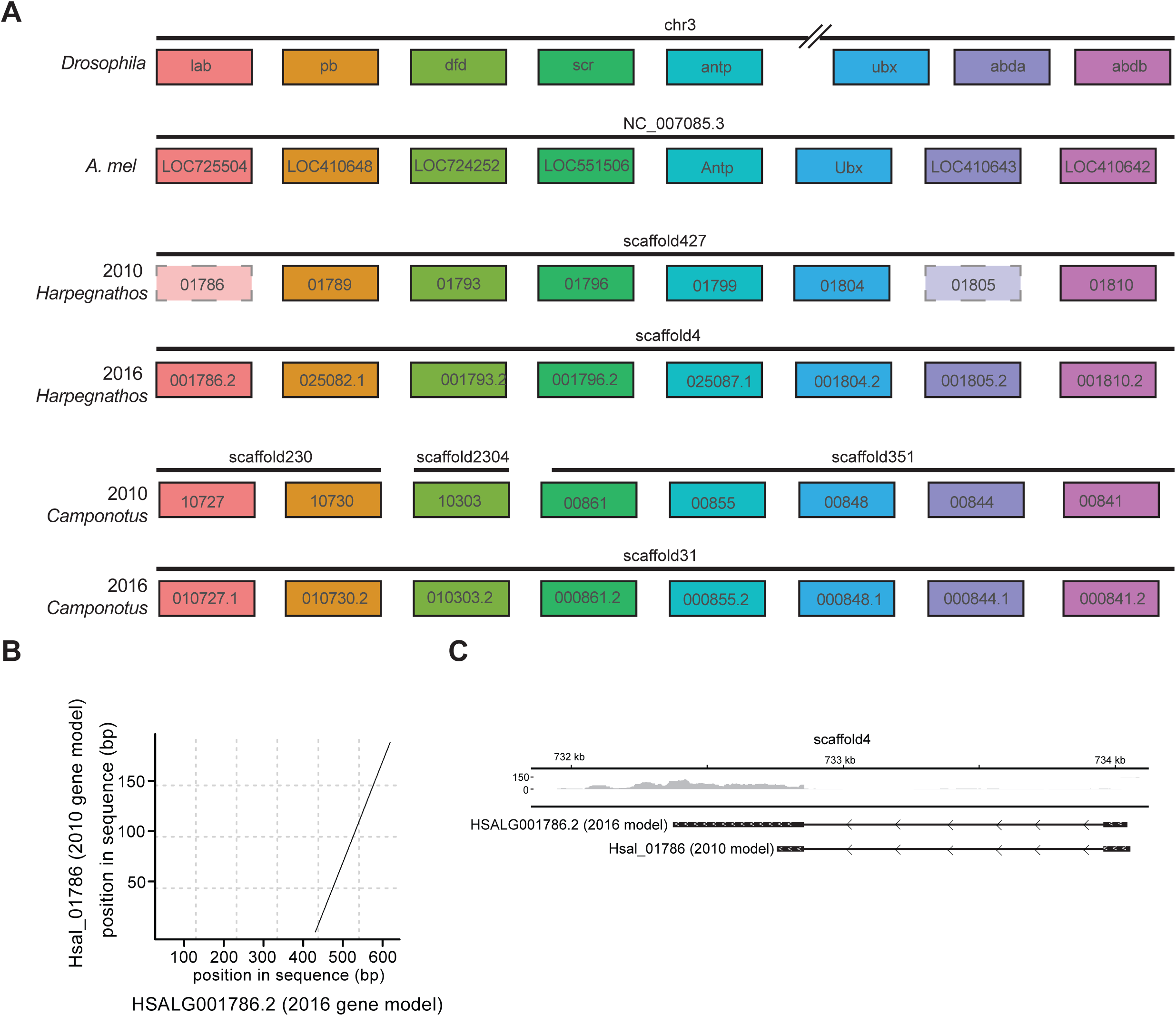
Reassembly of the *Hox* clusters of *Camponotus* and *Harpegnathos*. **(A)** The 2016 *Harpegnathos* annotation contains two *Hox* genes missing in the 2010 annotation (dashed boxes), and both the 2016 *Harpegnathos* and *Camponotus* annotations contain all *Hox* genes on the same scaffold, in contrast to the 2010 *Camponotus* annotation. **(B)** Example of a *Hox* gene in *Harpegnathos* updated in 2016 annotation. Hsal_01786 in 2010 annotation has homology to the corresponding gene model HSALG001786.2 in 2016 assembly, but only covers 33% of HSALG001786.2 and thus was not detected as a homolog of *Drosophila lab*. The 2010 gene model is depicted on the y-axis, with the 2016 gene model on the x-axis. Dots in the plot indicate regions of significant sequence similarity between 2010 and 2016 models. **(C)** RNA-seq from various developmental stages in *Harpegnathos* shows extension of the gene model past the 2010 boundaries. The 2010 and 2016 gene models are shown under the RNA-seq coverage track. Scale on RNA-seq track indicates reads per million.

Together, our analyses show that re-annotation of the improved 2016 genome assemblies for *Harpegnathos* and *Camponotus* yielded more complete gene sets, better models of already annotated genes, and, at least in one case, better contiguity of a tightly regulated gene cluster.

### Annotation of long non-coding RNAs in *Harpegnathos* and *Camponotus*

Having generated genome assemblies with greatly improved contiguity and protein annotations with more accurate gene models, we next sought to annotate lncRNAs. Toward this end, we performed a genome-guided *de novo* transcriptome assembly from RNA-seq of various developmental stages and detected high-confidence transcripts longer than 200 bp and not overlapping with existing protein-coding gene models (**Fig. S7**). About 24% of *de novo*-assembled *Harpegnathos* and *Camponotus* transcripts met this requirement (**Fig. 5A**).

**Figure 5.**
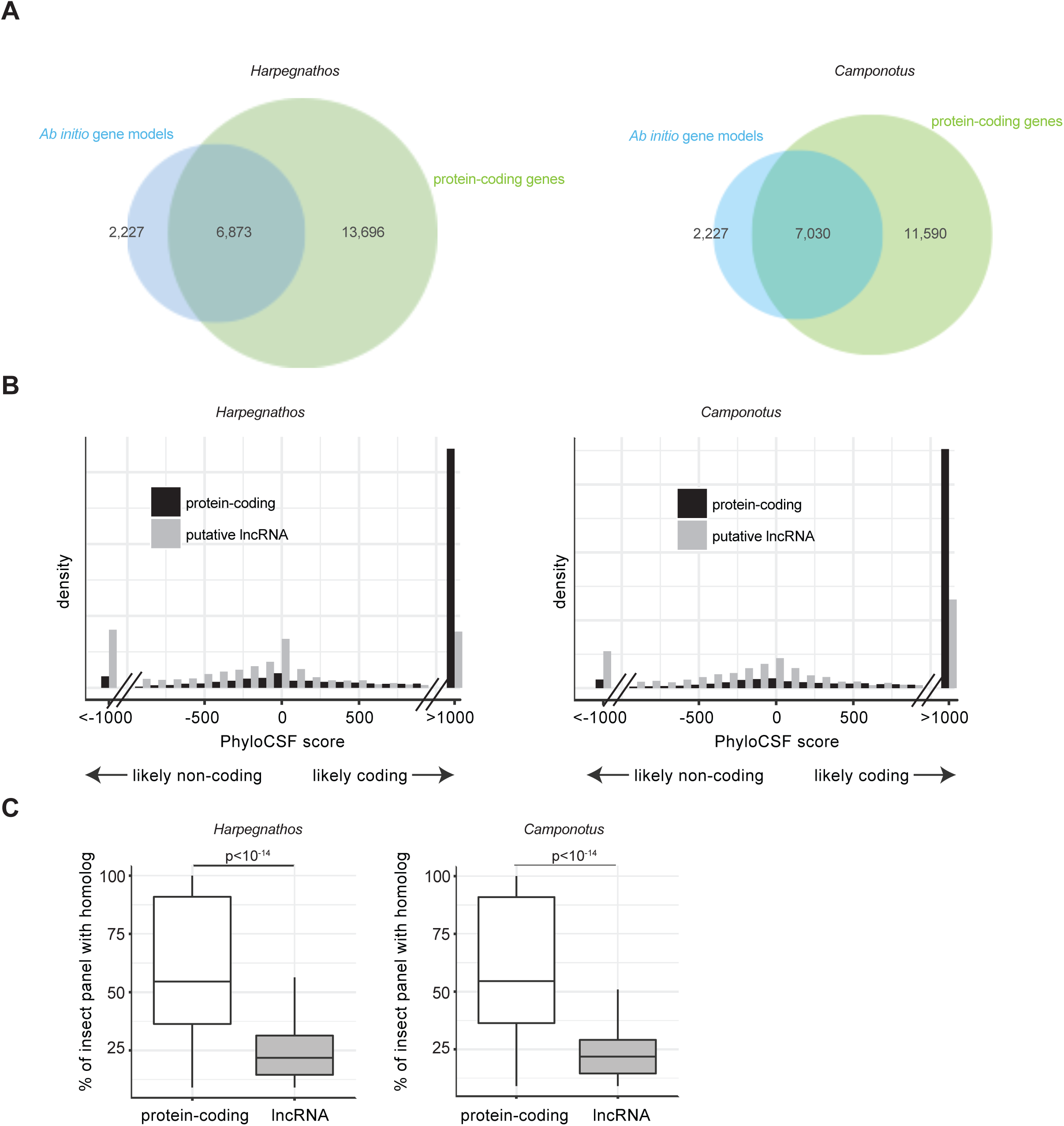
Annotation of long non-coding RNAs. **(A)** Venn diagram for the overlap between *ab initio* transcript assembled by Trinity and Stringtie with protein-coding gene models in *Harpegnathos* (left) and *Camponotus* (right). **(B)** PhyloCSF scores for transcripts with no overlap to coding sequences (gray) and known protein-coding genes (black). The x-axis indicates the PhyloCSF scores in decibans, which represent the likelihood ratio of a coding model vs. a non-coding model. Negative values indicate that a gene model is more likely to be coding than non-coding. **(C)** Boxplot for the number of homologs (BLASTN e-value < 10^-3^) found in other insect genomes for putative lncRNAs compared to protein-coding gene models. The transcriptomes of 54 insects and 1 outgroup (*Homo sapiens*) were used.

We further subdivided these putative non-coding transcripts into intervening, promoter-associated, and intronic, according to their spatial relationship with protein-coding annotations (**Fig. S8A**). To refine our non-coding annotations and confirm their lack of coding potential, we filtered them using their PhyloCSF score, a metric that takes into account the synonymous and non-synonymous mutation rate for any potential open reading frame (ORF) within a transcript by comparing it to homolog sequences across species^56^. A negative PhyloCSF score indicates that a transcript is not under evolutionary pressure to maintain a coding sequence and is therefore more likely to be non-coding. As expected, most protein-coding genes in both *Harpegnathos* and *Camponotus* had a high PhyloCSF score, whereas our newly annotated putative non-coding transcripts were skewed toward lower PhyloCSF scores (**Fig. 5B**), irrespective of their location relative to protein-coding genes (**Fig. S8B**).

**Figure 6.**
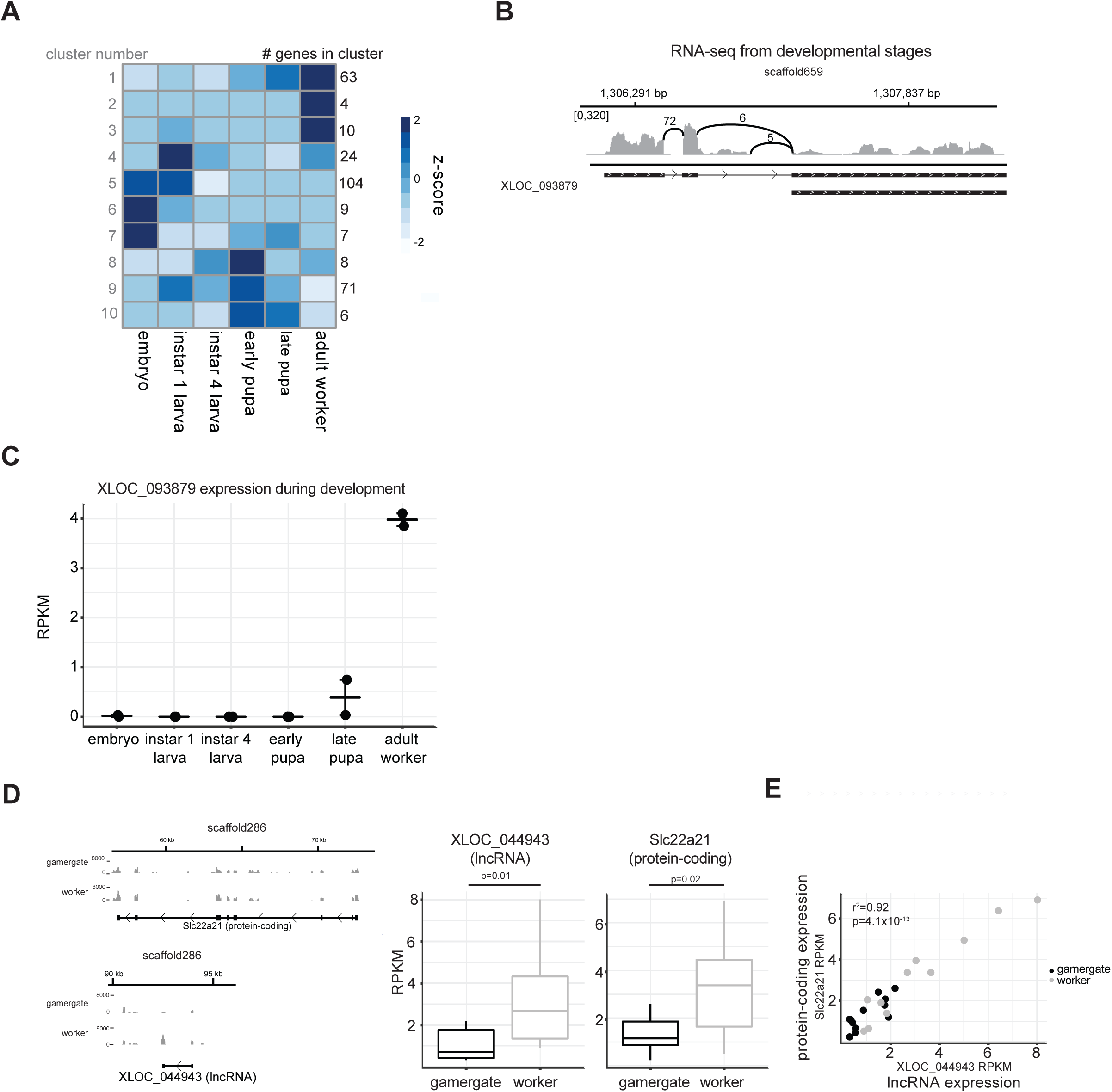
Differential expression of lncRNAs in *Harpegnathos* castes and developmental stages. **(A)** K-means clustering of changes in lncRNA expression shows characteristic expression patterns. Clustering was performed using all lncRNAs showing differential expression between any two developmental stages. The cluster number is displayed to the left of the heatmap, while the number of lncRNAs in each cluster is shown to the right. **(B)** Sashimi plot of RNA-seq from various developmental stages (see methods) covering an example lncRNA from cluster 2, XLOC_093879. The gene has three exons and two isoforms. Scale indicates number of reads **(C)** XLOC_093879 has expression levels typical of its cluster (**A**, cluster 2). Expression is low from the embryo stage until late papa, when it rises slightly. Adult workers display the highest expression. **(D)** A lncRNA, XLOC_044943, and its neighboring protein-coding gene, HSALG013780, are differentially expressed between brains from gamergates (n=12) and workers (n=11) in *Harpegnathos*. A genome browser snapshot (left) shows a pileup of reads on the exons of both genes, with higher peaks in worker brains. Scales on RNA-seq tracks indicate read per million. Quantification in RPKMs are shown to the right. P-values are from two-sided Student’s t-tests. **(E)** The expression levels of the lncRNA (x-axis) and the protein-coding gene (y-axis) shown in **(D)** correlate in both gamergate and worker. Each dot represents one biological sample (gamergate, n=12; worker, n=11). P-value from Pearson correlation is indicated.

For the final list of lncRNAs, we set a PhyloCSF score of −10 as threshold, which indicates that a given transcript is 10 times more likely to be non-coding than protein-coding. After filtering by PhyloCSF score and length, 628 (28.2%) and 683 (30.1%) of the putative non-coding transcripts in *Harpegnathos* and *Camponotus*, respectively, were designated lncRNAs. 15.5% *Harpegnatos* and 12.4% *Camponotus* protein-coding transcripts passed this threshold of −10 as well, suggesting an estimate of the false discovery rate for these predictions.

Previous efforts toward annotating lncRNAs have indicated a consensus set of features typical of lncRNAs in a variety of organisms: they are less conserved, shorter, have less exons, and, overall, are expressed at lower levels than protein-coding genes^57^. We detected most of these features in our ant lncRNAs: they were less conserved than protein-coding genes (**Fig. 5C**), regardless of their genomic localization (**Fig. S8D**); they had a smaller number of exons, with a majority of lncRNAs having only one (**Fig. S9A**); and they were typically expressed at lower levels than protein-coding genes, though not significantly in *Harpegnathos* (**Fig. S9B**). However, the length distribution of the ant lncRNAs was similar to that of protein-coding genes (**Fig. S9C**), which was a departure from what observed in mammals, *Drosophila*, and *C. elegans*^24,32,58,59^. As expected, virtually none of the lncRNAs had annotated PFAM domains, in contrast to protein-coding genes (**Fig. S9D**).

### Developmental and caste-specific lncRNAs

We next sought to determine whether lncRNA transcription was differentially regulated during major life transitions in *Harpegnathos*. First, we analyzed whole-body RNA-seq datasets from embryos, larvae, pupae, and adult workers. We clustered relative changes in the expression levels of lncRNAs across these samples into groups with distinct kinetics (**Fig. 6A**), which allowed us to identify early development lncRNAs (**Fig. 6A**, cluster 4, 5, 6, 7), adult lncRNAs (**Fig. 6A**, cluster 1, 2, 3), and a very interesting set of lncRNAs exclusively or predominantly expressed in the pupal stage (**Fig. 6A**, cluster 8, 9, 10) a critical phase in the life of holometabolous insects characterized by pronounced cell proliferation and morphogenesis. One illustrative example from the set of “adult lncRNAs” (cluster 2) is XLOC_093879, which gives rise to two isoforms containing one or three exons (**Fig. 6B**). The expression pattern was consistent with its cluster membership, with no expression in early developmental stages, low expression in late pupa, and high expression in adult workers (**Fig. 6C**, **Fig. S10A**). To confirm XLOC_093879 was a *bona fide* lncRNA, we used an orthogonal method of measuring coding potential used in many other lncRNA annotations^29,30,32,33^, the Coding Potential Calculator (CPC)^60^. CPC finds ORFs in a transcript sequence, compares ORF sequences to a database of protein-coding transcripts, and uses a support vector machine classifier to designate transcripts as coding or non-coding. Indeed, XLOC_093879 transcripts have CPC scores that classifies the gene as non-coding.

Finally, we wished to identify lncRNAs that might be differentially regulated during a behavioral switch. Using the same worker and gamergate RNA-seq described above (**Fig. 3C**), we analyzed changes in the expression of lncRNAs. We found 3 lncRNAs that were differentially expressed in worker and gamergate brains using a 10% FDR cutoff. An example from these, XLOC_044943, was also classified as a non-coding gene by its CPC score, and had higher expression in workers, similar to its neighboring protein-coding gene, *Slc22a21* (**Fig. 6D**). *Slc22a21* belongs to a family of organic solute transporters. Although *Slc22a21* itself has not been implicated in neuron function, other members of the family have been shown to function in the brain^61^. Interestingly, the expression levels of the protein-coding gene and the lncRNA correlated in multiple worker and gamergate brain samples (**Fig. 6E**), suggesting that the coding genes are co-regulated, or, possibly, that the lncRNA controls expression of the protein-coding gene, as in several cases of *cis*-acting lncRNAs in other organisms^62^. The lncRNA and protein-coding gene in this example are ∼20 kb apart on new scaffold286, but were assigned to different smaller scaffolds in the old annotation, which would have masked their potential for being co-regulated *in cis* (**Fig. S10B**). The correlation between the lncRNA and protein-coding gene expression was not due simply to the proximity of the genes, as other gene pairs at a similar or smaller distance did not display such strong correlation in expression (**Fig. S10C**).

Thus, our improved genome assemblies allowed us to generate a high-quality annotation of lncRNAs, several of which displayed developmental- and caste-specific expression patterns, and to uncover, in at least one case, a caste-specific lncRNA that might be involved in a *cis*-regulatory circuit in the brain.

## DISCUSSION

Social insects offer a unique perspective to studying epigenetics^1,2^. Striking morphological and behavioral differences between castes include phenotypes relevant to translational research, such as social behavior, aging, and development. These traits can be studied on an organism level within a natural social context, as full colonies can be maintained in the laboratory. However, to analyze these complex traits at a molecular level, proper genomic tools must be developed. We previously assembled the first ant genomes generating a workable draft using the best technology at the time: whole-genome shotgun using short Illumina reads^5^. Although the release of the *Camponotus* and *Harpegnathos* genomes, along with additional ant genomes following shortly after, spurred a number of studies on ant genomics and epigenomics, the draft quality of the genomes remained an obstacle to more sophisticated analyses.

Moving a genome beyond draft status often entails connecting disjointed contigs using mate pair scaffolding^63^, proximity ligation technologies such as Hi-C^64–66^, or optical mapping with restriction enzymes^67^. Recently, long reads from third-generation sequencing technologies have been used to scaffold short-read assemblies^35^,or, with sufficient sequencing depth, to construct a *de novo* assembly. A few eukaryotic genomes have been assembled *de novo* with long reads^67–70^ or by using long reads as scaffolds^71^, but our assembly is the first social insect genome to be assembled using this strategy. One major advantage granted by long reads is the improved ability to assemble over repeats, which typically cannot be resolved with short reads^39^, generally improving genome contiguity.

Here, we used PacBio long reads to reassemble *de novo* the genomes of the two ant species currently in use as molecular models in our laboratory, *Camponotus floridanus* and *Harpegnathos saltator*. These new assemblies reached scaffold N50 sizes larger than 1 Mb (**Table 1**) while resulting in increased sequence accuracy as measured by RNA-seq mapping rates and mismatches and comparison with Sanger sequencing of fosmid clones (**Fig. 2**). Perhaps more importantly, the number of gaps and gapped bases disrupting the continuity of the new scaffolds is smaller than in all other insect genomes available on NCBI (**Fig. 1E**). These greatly improved assemblies deliver several critical benefits that are indispensable to further develop these ant species into molecular model organisms: 1) more comprehensive protein-coding annotations and more complete gene models (**Fig. 3, Table 2**); 2) more continuity of co-regulated gene clusters (**Fig. 4**); 3) the ability to annotate lncRNAs with confidence (**Fig. 5**), and 4) the ability to detect regulatory mechanisms functioning in *cis* at large genomic distances (**Fig. 6**). We discuss the implications of some of these points more in detail below.

Although the annotation of protein-coding genes did not suffer excessively from the draft status of the 2010 assemblies, the new annotations contain potentially relevant genes that were previously missing. Most notably, a *Gp-9-like* gene was newly annotated in the *Harpegnathos* genome and found to be differentially expressed in worker brains compared to gamergates (**Fig. 3C**). The importance of *Gp-9* in ant biology is well established as it was one of the first genetic markers discovered in ants for a colony-level phenotype, the choice between a polygyne (multiple queens) or monogyne (single queen) social form in colonies of the fire ant *Solenopsis invicta*^72^. Recent genomic studies revealed that *Gp-9* maps to a cluster of genes involved in a large genomic rearrangement that distinguishes the two social forms and has given rise to a so-called “social chromosome”^45^. However, the role of *Gp-9* in social behavior remains unknown.

Our new finding that a *Gp-9-like* homolog is expressed at different levels in *Harpegnathos* castes led us to a reanalysis of expression patterns in this gene family in two other ant species, *Dinoponera quadriceps* and *Vollenhovia emeryi*. *Dinoponera* colonies are queenless, instead comprising non-reproductive workers (“beta” or “low”) and one ant (“gamergate” or “alpha”) that lays eggs. The top ranking worker can replace the reproductive upon death or removal of the alpha^73^, similar to *Harpegnathos* workers gaining the license to reproduce upon isolation from the queen. *Vollenhovia* colonies also have an unusual social structure, in which queens can produce a clonal queen via thelytoky, as well as workers and sexual queens via sexual reproduction, and males can be produced through androgenesis^74^. In each of these ants, at least one *Gp-9-like* homolog was expressed at significantly higher levels in the non-reproductive phenotype compared to the reproductive phenotype. The finding that *Gp-9*, already implicated in contributing to the unusual colony structure of *Solenopsis invicta*, has homologs that are expressed at different levels in the castes of three additional ant species opens an avenue for future investigation on its molecular function.

Another key advance granted by our improved genome assemblies was the ability to annotate lncRNAs with confidence. We developed a custom pipeline and discovered over 600 lncRNAs with very low coding potential according to evolutionary analysis of their sequence by PhyloCSF^56^ in both *Harpegnathos* and *Camponotus*. The mechanism of action and biological impact of lncRNAs is the subject of intense investigation in various model systems and in several cases a dedicated role in brain function has been advocated, based in part on their expression patterns^1,13^. Although a few cases of *trans* regulatory activity for lncRNAs have been demonstrated, it is generally believed that lncRNAs act in *cis* to regulate expression of neighboring genes^13,62,75^. Therefore, an extended view of protein-coding genes in the vicinity of lncRNAs is critical to construct hypothesis on their regulatory role, and this information is provided by our updated genomes. Indeed, thanks to the increased continuity of the new assemblies, we were able to identify a lncRNA–mRNA pair whose brain expression levels were co-regulated and different between *Harpegnathos* workers and gamergates. The fact that the mRNA in this example encodes a protein from a family of membrane channels involved in brain function^61^ further suggests that this regulatory interaction might be important for caste-specific behavior.

The improved genome assemblies of *Camponotus* and *Harpegnathos* ants will also be instrumental in the analysis of enhancer-mediated regulation of gene expression in different castes. A growing amount of evidence suggests that enhancers rather than promoters are key to understand how genomes encode organismal complexity^76,77^. However, enhancers can act at considerable genomic distances^78^, regulating gene expression by coming into contact with promoters via chromatin looping. Therefore, a fragmented genome assembly would likely place enhancers in a scaffold different from that containing the gene they regulate, preventing correct analysis of their function. Genome-wide patters of histone H3 lysine 27 acetylation, a chromatin mark typically associated with enhancer function, are strongly predictive of caste identity in *Camponotus floridanus*^7^, and artificial changes in its levels are sufficient to stimulate caste-specific behavior^8^. The improved assemblies will facilitate further molecular dissection of this phenomenon.

Finally, the benefits of an upgraded genome go beyond gene model annotation and *cis* regulatory elements. Transposable elements exist in insects and play a major role in the evolution of insect genomes^79,80^. Our new assemblies capture more repeat content, including a large amount of species specific repeats, and perhaps will contribute to the growing understanding of genome evolution and structure in insects.

## EXPERIMENTAL PROCEDURES

### Ant colonies and husbandry

Ants were housed in plaster nests in a clean, temperature- (25°C) and humidity-(50%) controlled ant facility on a 12-hour light/dark cycle. *Harpegnathos* ants were fed three times per week with live crickets. *Camponotus* ants were fed twice weekly with excess supplies of water, 20% sugar water (sucrose cane sugar), and Bhatkar-Whitcomb diet^81^. The *Harpegnathos* colony was descended from the colony sequenced for the original 2010 genome assembly, which was originally collected as a gamergate colony in Karnataka, India in 1999 and bred in various laboratories since^4,5^. The *Camponotus* colony was collected in Long Key, Florida in November 2011.

### Long read DNA library preparation and sequencing

High molecular weight genomic DNA was extracted from 36 *Harpegnathos* and 42 *Camponotus* recently eclosed workers. Gasters were removed before sample homogenization to reduce contamination from commensal bacteria. Size selection and sequencing was performed by the University of Washington PacBio Sequencing service using BluePippin size selection and P6-C4 chemistry, RSII platform. Reads of insert (ROIs) were extracted using SMRT analysis software. The RS_ReadsOfInsert.1 protocol was used, with the parameters 0 minimum full passes and 75% minimum predicted accuracy. 34 SMRT cells were processed for *Harpegnathos*, producing 3.1x10^6^ ROIs containing 2.3x10^10^ total bases, for a mean ROI length of 7,471 bp. 17 SMRT cells were processed for *Camponotus*, producing 1.1x10^6^ ROIs containing 1.0x10^10^ total bases, for a mean ROI length of 9,934 bp.

### Repeat masking and evaluation of repeats in new sequence content

Although repeat masking was performed by the MAKER2 pipeline internally during the protein-coding gene annotation step, RepeatMasker (A.F.A. Smit, R. Hubley & P. Green RepeatMasker at http://repeatmasker.org) was also run independently to compare repeats in the 2010 genome assemblies to the 2016 assemblies and to produce a masked genome FASTA. First, the genomes were masked with RepeatMasker and the “*Harpegnathos saltator*” library. Custom repeat libraries were then constructed using RepeatScout on the 2016 genomes with default parameters. These libraries were used in RepeatMasker to find species-specific repeats. Next, we detected non-interspersed repeat sequences with RepeatMasker run with the “-no int” option. Finally, we used Tandem Repeat Finder^82^ with the following parameters: match=2, mismatch=7, delta=7, PM=80, PI=10, minscore=50, MaxPeriod=12.

To detect new sequence content, the 2010 genomes were broken into 500 bp non-overlapping windows, then aligned to the 2016 assemblies using Bowtie2^83^.

### Genome assembly strategy

The extracted ROIs were error corrected, trimmed, and assembled by Canu v1.3^34^. Error correction and assembly were performed with default parameters with the following changes: corMhapSensitivity = high, corMinCoverage = 0, errorRate = 0.03, minOverlapLength = 499. Quiver was used to polish the assemblies, using the SMRT Analysis protocol RS_Resequencing with default parameters. Scaffolding using both long reads and mate pairs was performed for both *Harpegnathos* and *Camponotus* assemblies, but mate pair scaffolding was done first in *Harpegnathos* and long read scaffolding was done first in *Camponotus*. SSpace-Standard^36^ was used to scaffold the assemblies using mate pair sequencing data with inserts of 2.2 kb (*Harpegnathos*: 5 libraries, *Camponotus*: 1 library), 2.3 kb (*Camponotus*: 1 library), 2.4 kb (*Camponotus*: 1 library), 2.5kb (*Harpegnathos*: 1 library), 5kb (*Harpegnathos:* 4 libraries, *Camponotus*: 2 libraries), 9kb (*Harpegnathos*: 1 library), 10kb (*Harpegnathos*: 1 library, *Camponotus*: 1 library), 20kb (*Harpegnathos*: 1 library, *Camponotus*: 1 library), or 40k (*Harpegnathos*: 1 library, *Camponotus*: 1 library). Standard parameters were used. For scaffolding with long reads, subreads were extracted from PacBio sequencing data using bash5tools with the following parameters: minLength=500, minReadScore=0.8. PBJelly^35^ was then used to perform the scaffolding, following the normal protocol. After scaffolding with mate pairs and PacBio subreads, the assemblies were polished using paired-end Illumina short reads and the tool Pilon to produce the final assemblies.

### Short-read DNA sequencing

Short-read DNA sequencing data (SRP002786)^5^ were used to polish the genome assemblies with Pilon. Reads were mapped to the *Harpegnathos* or *Camponotus* genome using Bowtie2 with default parameters. Due to memory limitations, the short DNA reads were aligned to the genomes in three sets. After the first set was used to polish the genomes, the reads from the next set were aligned to the consensus sequence produced using the previous set.

### Comparison of 2016 *Harpegnathos* and *Camponotus* assemblies to other insects

Other insects used for comparison included all insects with scaffold-level genomes annotated by NCBI as of 5/8/17 (n=81). Scaffold number, contig number, scaffold N50, contig N50, number of gaps, and number of gapped bases were obtained from the genome FASTA available for download on the NCBI website.

BLAST was used to find homologs to *Harpegnathos* and *Camponotus* genes in the 2010 and 2016 annotations. We searched an ant panel consisting of 16 ants (*Wasmannia auropunctata, Pogonomyrmex barbatus, Cerapachys biroi, Atta cephalotes, Atta colombica, Trachymyrmex cornetzi, Cyphomyrmex costatus*, *Acromyrmex echinatior, Vollenhovia emeryi, Linepithema humile, Solenopsis invicta, Monomorium pharaonis, Dinoponera quadricepts, Trachymyrmex septentrionalis, Trachymyrmex zeteki*) and a Hymenoptera panel consisting of 16 non-ant Hymenopterans (*Orussus abietinus, Diachasma alloeum, Ceratina calcarata, Polistes canadensis, Apis cerana, Microplitis demolitor, Polistes dominula, Apis dorsata, Apis florea, Copidosoma floidanum, Bombus impatiens, Trichogramma pretiosum, Megachile rotunda, Bombus terrestris, Nasonia vitripennis*). To qualify for “all insects” in **Fig. 3A**, the gene had to have a homolog in at least 90% of ants, Hymenoptera, and in *Drosophila melanogaster*. To qualify for “mammals and insects,” the gene had to meet the same requirements for “all insects” and have a homolog in both *Mus musculus* and *Homo sapiens*.

### Fosmid analysis

Ten Sanger sequenced fosmids^5^ with an average length of 36,755 bp were analyzed for *Harpegnathos,* and 11 fosmids with a mean length of 37,610 bp were analyzed in *Camponotus*. The scaffold with the most hits for each fosmid in both 2010 and 2016 genome assemblies was found using BLAST. Next, the fosmid and the scaffold with the closest matches were globally aligned. The coverage (how many of the fosmid bases matched with the genome) and the length of the scaffold containing the fosmid were reported.

### Developmental stage RNA sequencing and analysis

RNA was extracted from the whole bodies of ants at various developmental stages for *Harpegnathos* (embryo, instar 1 larva, instar 4 larva, early pupa, late, pupa, adult worker, adult male) and *Camponotus* (embryo, instar 1 larva, instar 4 larva, late pupa minor, late pupa major, minor worker, male). For library preparation, 500 ng polyA+ RNA was isolated using Dynabeads Oligo(dT)_25_ (Thermo Fisher) beads and constructed into strand-specific libraries using the dUTP method^84^. UTP-marked cDNA was end-repaired (Enzymatics, MA), tailed with deoxyadenine using Klenow exo^-^ (Enzymatics), and ligated to custom dual-indexed adapters with T4 DNA ligase (Enzymatics). Libraries were size-selected with SPRIselect beads (Beckman Coulter, CA) and quantified by qPCR before and after amplification. Sequencing was performed on a NextSeq 500 (Illumina, CA) in a 200/100 bp paired end format. The 200 bp were aligned to the genome using STAR^85^ with default parameters, but after clipping 75 bp from the 3’ end due to decreasing sequence quality. The mapping rate and mismatch rate per base were reported by STAR. Read counts were calculated for each gene or lncRNA using HTSeq-count^86^.

### Annotation of protein-coding RNAs

Protein-coding genes were annotated on the *Harpegnathos* and *Camponotus* assemblies using iterations of the MAKER2 pipeline^41^. Inputs to the protein homology evidence section of MAKER2 were FASTA files of proteins in *Apis mellifera, Drosophila melanogaster,* and the previous *Harpegnathos* or *Camponotus* annotation. RNA-seq was provided as EST evidence. RNA-seq was processed using PASA_Lite, a version of PASA^87^ that does not require MySQL. First, a genome guided transcriptome reassembly was produced using Trinity^88^. The transcriptome was aligned against the genome using BLAT with the following parameters: -f 3 –B 5 –t 4. The alignments were used as input to PASA_Lite, which produces spliced gene models. The PASA_Lite output was further processed with TransDecoder^89^, a tool that searches for coding regions within transcripts.

The first iteration of MAKER2 was run with the settings est2genome=1 and protein2genome=1, indicating that both models directly from RNA-seq and homology mapping were output. No SNAP^90^ hidden Markov model (HMM) was provided in the first iteration. Augustus^91^ HMMs were provided; in the first run of maker, the *Camponotus*_*floridanus* parameters provided with Augustus were used for *Camponotus*, and parameters trained on an earlier version of the *Harpegnathos* genome were used for *Harpegnathos*. After the first MAKER2 run, SNAP and Augustus HMMs were trained using the output of the previous step. High confidence gene models were extracted using BUSCO v2^44^, a tool that measures the completeness of a transcriptome set. BUSCO searches for the presence of conserved orthologs in the transcriptome, and also can produce a list of which genes are complete gene models. Only these complete models were used to train Augustus and SNAP.

The second iteration of MAKER2 was run with the same homology and RNA-seq inputs, but with the new HMMs and the GFF from the previous step included as an option in the Re-annotation parameters section, and with est2genome=0 and protein2genome=0. After the second MAKER2 iteration, HMMs were trained using the same steps as above, and the process was repeated two more times. On the fourth MAKER2 run, est2genome and protein2genome were turned on, producing gene models directly from RNA-seq and homology. The gene models from the last iteration of MAKER2 were filtered using the reported annotation edit distance (AED, measures the level of agreement between different sources of evidence) and the presence of a PFAM domain. PFAM domains were detected using HMMer v3.1b2 (http://hmmer.org) with the PFAM-A database. Genes were retained if they had either an AED < 1 or a PFAM domain, or both.

Gene identifiers (IDs, e.g. HSALG000001) were assigned to genes based on the presence of homolog in the 2010 annotation. If the 2016 had a perfect match at the nucleotide level in the 2010 assembly, it retained the old ID with the version 1 (e.g. HSALG000001.1). If the 2016 model significantly matched at the protein level, but not at the nucleotide level, it retained the old ID with the version 2 (e.g. HSALG000001.2). If multiple 2010 genes were significant matches, multiple 2016 genes matched to the same 2010 gene, or no homolog was present in the old assembly, a new ID was issued.

### Assessment of annotation quality

The transcriptome completeness was measured using BUSCO v2, which searches for the presence of well conserved orthologs in a transcriptome. The *arthopoda* set was used as the test lineage.

### *Hox* cluster analysis

To detect whether the genome annotation captured the genes in the *Hox* cluster, we searched for *Drosophila melanogaster Hox* genes in the *Apis mellifera* genome, as well as the 2010 and 2016 *Harpegnathos* and *Camponotus* annotations. The gene was denoted as present if there was a significant (e-value < 10^-5^) hit using standard megablast parameters.

### *Gp-9* homologs differential expression

RNA-seq from full bodies of *Vollenhovia emeryi* (PRJDB3517, RNA-seq from 5 queens and 5 workers)^74^ and *Dinoponera quadriceps* (GSE59525, RNA-seq from 7 alpha and 6 low ants)^92^ was aligned to the genome and mapped to NCBI annotated features. All genes annotated as “*Gp9*” or a “*Gp9-like*” were evaluated for differences in expression between reproductive (queen or alpha) and non-reproductive (worker or low) ants. RPKMs between castes were compared using Student’s t-tests.

### Annotation of lncRNAs

RNA-seq reads from various developmental stages of *Harpegnathos* (embryo, instar 1 larva, instar 4 larva, early pupa, late pupa, adult worker, male) and *Camponotus* (embryo, instar 1 larva, instar 4 larva, late pupa minor, late pupa major, minor, male) were assembled using two genome-guided *de novo* assemblers, Trinity^87^ and Stringtie^93^. The transcripts produced from these two methods were merged using cuffmerge^94^, then each reassembled transcriptome was intersected (reciprocal 75% overlap required) with the merged transcripts to produce a file for each method with transcripts from the same set. Transcripts from both methods were then intersected (required 75% reciprocal overlap). Finally, this high-confidence transcriptome was intersected with the coding sequences of protein-coding genes, and only transcripts with no overlap to protein-coding genes were designated as intergenic. Transcripts were further split by location for some analyses: “intervening” denotes no overlap with protein-coding genes, “intronic-sense” indicates the transcript is an intron of a gene in the same orientation, “intronic-antisense” indicates the transcript is in an intron of a gene in the opposite orientation, “intronic-both” indicates the gene is intronic to a gene in the sense and antisense direction, and “promoter-associated” indicates that the lncRNA overlaps is within 1,000 bp of a promoter of an antisense gene. The intergenic transcripts were collapsed into loci based on cuffmerge results for some analyses.

BLAST was used to find homologs for intergenic transcripts in a panel of 54 insects and an outgroup (human). Only hits with an e-value of ^10–3^ were kept. A multispecies alignment was performed for each transcript using MAFFT. TimeTree^95^ was used to create a phylogeny complete with branch lengths of the insect panel and either *Harpegnathos* or *Camponotus*. The phylogeny was rooted using the R package *ape*, with *Homo sapiens* as the outgroup. Using this phylogeny and the multispecies alignment, the PhyloCSF Omega Test mode was run, with all reading frames in the sense direction tested, to assess the coding potential of each transcript. PhyloCSF scores are given in the form of a likelihood ratio, in the units of decibans. A score of x means the coding model is x times more likely than the non-coding model (for example, if x=10, the coding model is 10 times more likely; if x=−10, the non-coding model is 10 times more likely). Transcripts with a score < −10 were considered lncRNAs.

Coding Potential Calculator (CPC)^60^ was used to confirm the non-coding status of lncRNA chosen as examples in the differential expression analyses. The UniRef90 database was used as a BLAST database.

### Clustering of lncRNA expression levels

Expression patterns of differentially expressed lncRNAs in the developmental stages of *Harpegnathos* (embryo, instar 1 larva, instar 4 larva, early pupa, late pupa, adult worker) were clustered using a quantile normalization of the log-fold expression (RPKM) change between each pair of samples. K-means clustering with a preset number of clusters (10) and maximum number of iterations (50) was performed on this quantile-normalized matrix. The heatmap of expression patterns was created using pheatmap, with color scaling by row.

### Sequencing data

RNA sequencing data generated for this study have been deposited in the NCBI GEO as SuperSeries GSE102605. PacBio reads and assemblies are being submitted to NCBI. Data will remain private during peer review and released upon publication.

## AKNOWLEDGMENTS

The authors thank J. Gospocic for providing ant samples and the UW PacBio Sequencing service for performing SMRT sequencing. R.B. thanks Danny Reinberg (NYU) as well as Guojie Zhang, Cai Li, Zhensheng Chen, and Luohao Xu (BGI) for their intellectual support and efforts during an initial attempt at annotating lncRNAs. R.B. acknowledges financial support from the NIH (DP2MH107055), the Searle Scholars Program (15-SSP-102), the March of Dimes Foundation (1-FY-15-344), a Linda Pechenik Montague Investigator Award, and the Charles E. Kaufman Foundation (KA2016-85223). E.S. acknowledges financial support from the NIH (T32HG000046). The authors thank Yoseph Barash, Ben Voight, and Paul Babb for comments on the manuscript.

